# Pathogenic *Rickettsia* species evade autophagosomal maturation and reduce anti-microbicidal pro-inflammatory IL-1 responses to support their intracellular survival

**DOI:** 10.1101/2022.06.19.496761

**Authors:** Oliver H. Voss, Hodalis Gaytan, M. Sayeedur Rahman, Abdu F. Azad

## Abstract

Species of genus *Rickettsia* are obligate intracellular bacterial parasites of a wide range of arthropod and vertebrate hosts. Some *Rickettsia* species are responsible for several serious human diseases. One fascinating feature of these stealthy group of pathogens is their ability to exploit host cytosolic defense responses to their benefits. However, the precise mechanism by which pathogenic *Rickettsia* elude host immune defense responses remains to be determined. Here, we observed that pathogenic *R. typhi* and *R. rickettsii*, but not non-pathogenic *R. montanensis* become ubiquitinated and induce autophagy upon entry into bone-marrow-derived macrophages. Moreover, unlike *R. montanensis, R. typhi*, and *R. rickettsii* colocalized with LC3B but not with Lamp2 upon host cell entry. Finally, we observed that pathogenic but not non-pathogenic *Rickettsia* reduce pro-inflammatory IL-1 responses. In sum, we identified a previously unappreciated pathway by which pathogenic, but not non-pathogenic, *Rickettsia* become ubiquitinated and induced autophagy, but avoided autophagolysosomal destruction as well as inflammasome-mediated anti-microbicidal IL-1 cytokine responses to establish an intracytosolic niche in macrophages.

## Introduction

A plethora of intracytosolic bacteria utilize sophisticated strategies to circumvent host defense mechanisms to promote their survival within the host (1,2). *Rickettsia* are arthropod-borne obligate intracellular bacteria with both symbiotic and pathogenic lifestyles. *Rickettsia* species (spp), such as *R. prowazekii, R. typhi, R. rickettsii*, and *R. conorii*, etiological agents for epidemic typhus, murine typhus, Rocky Mountain spotted fever, and Boutonneuse Fever, respectively; are among the most virulent pathogens (3,4). Apart from their historical record, the global impact of rickettsial infections is illustrated by the resurgence of known, as well as the rise of unrecognized pathogens (5). Infections of humans with *R. rickettsii* continue to cause severe consequences in South and Central America (6), and the resurgence of *R. conorii* in Europe, the Middle East, and Africa highlights current threats of rickettsial diseases (7). In addition, arthropod-borne rickettsial diseases are also on the rise in the USA, as exemplified by recent outbreaks of *R. rickettsii* in Arizona (8) and of *R. typhi* in California (9) and Texas (10).

*Rickettsia* entry into host cells, followed by intracytosolic growth and dissemination to neighboring cells has long been considered as a conserved process. However, it is important to note (and often overlooked) that many described rickettsiae are naturally maintained in arthropods and are considered non-pathogenic, as they cause no or a very mild disease in humans (3,11,12). *Rickettsia* spp are introduced into the host’s dermis by infected arthropods and the first host defense cells encountered are macrophages and dendritic cells (4). In particular, macrophages are crucial in either terminating an infection at an early stage or succumbing to pathogen colonization, thereby contributing to the dissemination of *Rickettsia* to distant organs of the host (4). Upon cell entry, *Rickettsia*, encounter cytosolic host defense responses initiated after sensing the bacteria or associated danger signals. Commonly invading bacteria will encounter autophagy and inflammasome responses, two functionally interconnected pathways (13,14), which typically provide the appropriate defense measures against intracellular pathogens (15,16). Importantly, recent reports suggest that autophagy acts on intracellular microbes upstream of the inflammasome (17,18). Autophagy is triggered by ubiquitination of the intracellular bacteria (19). Ubiquitin (Ub) coats the bacterial surface and recruits autophagy adaptors (p62, NDP52) and induces autophagosome formation with autophagy machinery consisting of ULK complex, PI3K complex (Beclin-1, VSP34, ATG14L) and the ATG16L1 complex (ATG16L1, ATG5, LC3) (20). It is becoming increasingly evident that several intracellular bacteria develop a unique mechanism to modulate autophagy, in particular to avoid autophagosomal destruction, to facilitate host colonization (15,19). In the case of *Rickettsia*, however, the role of autophagy in regulating host colonization remains inconclusive (21–24). For instance, *R. australis*, a virulent member of the transitional group (TRG), benefit from ATG5-dependent autophagy induction and suppression of pro-inflammatory cytokine responses to colonize host cells (21,22). In contrast, reports on *R. parkeri*, a less-virulent member of the spotted fever group (SFG), demonstrated that its surface protein OmpB is critical for protecting against autophagic recognition, while evasion of autophagy was critical for rickettsial invasion of BMDMΦ and WT mice (23,24). Intriguingly, we reported that pathogenic members of SFG and typhus group (TG) *Rickettsia* secret effectors to promote host colonization by modulating endoplasmic reticulum structures or by hijacking the autophagic defense pathway (25–30). In fact, our findings on the flea-transmitted *R. typhi* (pathogenic member of TG), showed that *R. typhi* is ubiquitinated upon host entry and escapes autolysosomal fusion to establish an intracytosolic niche in non-phagocytic cells (29). Thus, in agreement with reports on *R. australis* (21,22), these findings suggest a mechanism by which highly virulent *Rickettsia* spp, including members of TG, TRG, and likely highly pathogenic SFG, activate autophagy, but subsequently evade autolysosomal destruction, to promote host colonization. These unexpected findings on how SFG, TRG, and TG *Rickettsia* differentially promote their host dissemination prompted us to test the hypothesis that pathogenic, but not non-pathogenic, *Rickettsia* spp utilize autophagy for evading lysosomal destruction and reducing inflammasome-mediated anti-microbicidal pro-inflammatory IL-1 responses to establish a replication niche in phagocytic host immune defense cells, like macrophages.

## Results

### Pathogenic *Rickettsia* are ubiquitinated and induce autophagy upon entry into macrophages

Preceding findings suggest that intracellular pathogens, like *Rickettsia*, not only encounter inflammasome-dependent defense mechanisms but are also confronted by another cytosolic defense pathway, autophagy (21–24,29,31–34). Both responses not only are key to mount the appropriate host defense responses, but are also functionally interconnected (15,16). In one of our preceding reports, we showed that *R. typhi* is ubiquitinated upon host entry, induces autophagy, but escapes autophagolysosomal maturation for intracellular colonization in non-phagocytic cells (29). More recently, we demonstrated that unlike *R. montanensis*, a non-pathogenic SFG member; *R. rickettsii* and *R. typhi*, two highly pathogenic members representing the SFG and TG, respectively, preferentially targeted the non-canonical inflammasome-IL-1α signaling axis in macrophages to supporting their replication. Given these reports by others and our recent findings, we first evaluated the ubiquitination status of *R. typhi, R. rickettsii*, or *R. montanensis*, during invasion of bone marrow-derived macrophages (BMDMΦ) isolated from C57BL/6J wild-type (WT) mice. Similar to infection studies using *R. australis* (22), we observed that both pathogenic *R. rickettsii*, and *R. typhi* strains were ubiquitinated upon entry into WT BMDMΦ (**Figures 1A, B**). In contrast, but similar to *R. parkeri* infections, *R. montanensis* was not ubiquitinated in WT BMDMΦ (**Figures 1A, B**), suggesting that non-pathogenic *R. montanensis* is likely destroyed by phagolysosomal fusion before escaping into cytosol of macrophages. Of note, our microscopy studies identified the utilized *Rickettsia* spp in different shapes and sizes, representing pleomorphic changes of *Rickettsia* during their intracellular growth cycle in the host cell (35,36). Furthermore, we evaluated the status of autophagy markers p62 (SQSTM1) and autophagic vesicle formation marker LC3B (37) during infection of WT BMDMΦ by Western blot analyses. Our data revealed that, unlike *R. montanensis*, infection with both pathogenic *Rickettsia* spp increased autophagic flux, as evidenced by an enhanced induction of LC3B and the simultaneously down-regulation of p62 expression (**Figures 1C, D**).

**Figure 1.**
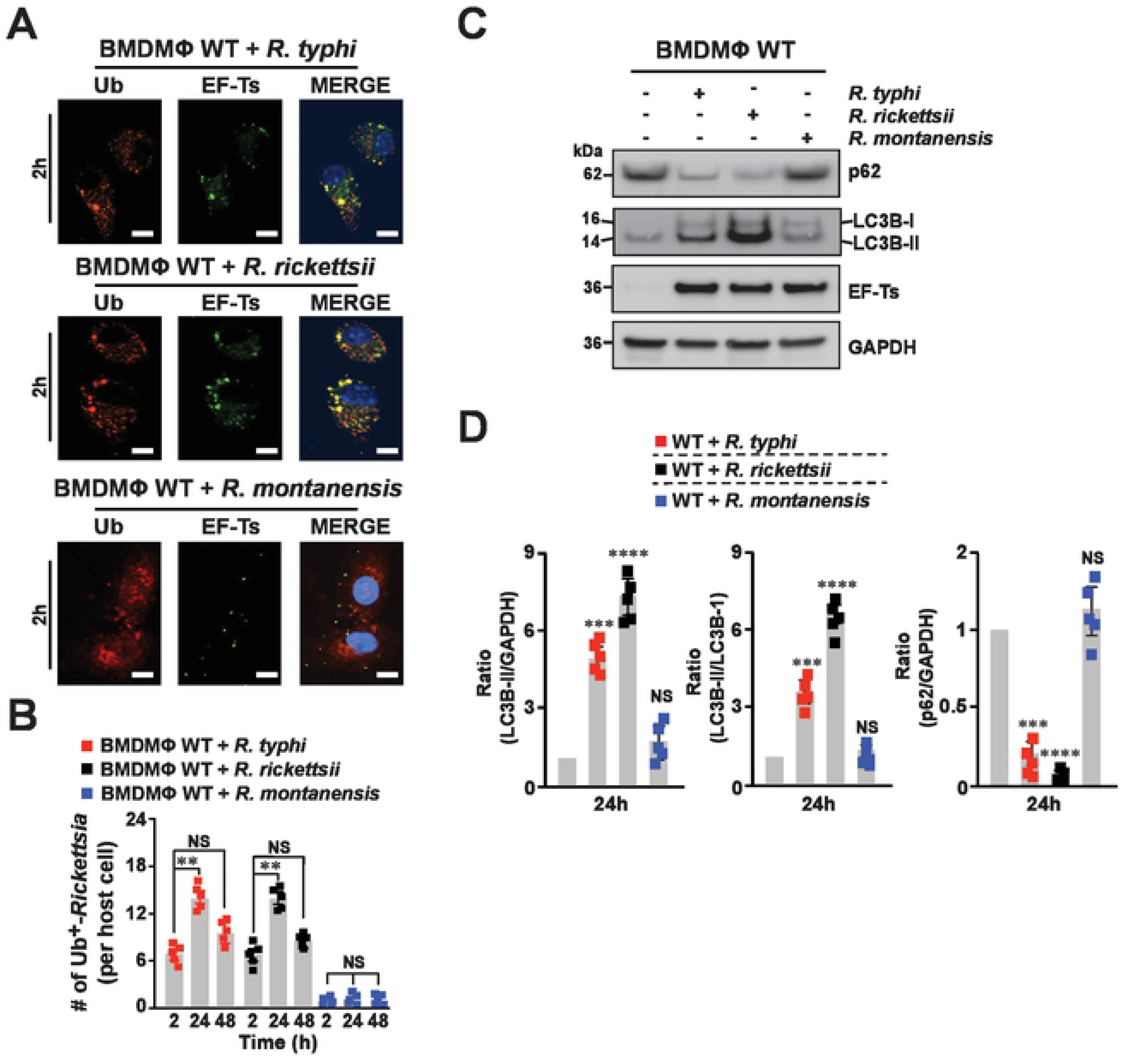
Pathogenic, but not non-pathogenic, *Rickettsia* species are ubiquitinated and induced autophagy in macrophages. (**A**) BMDMΦ from WT mice were infected with *R. montanensis*, *R. rickettsii*, or *R. typhi*. Samples were fixed with 4% PFA and *Rickettsia* spp were detected using the rickettsial specific Alexa Fluor 488-conjugated EF-Ts antibody, while ubiquitination (Ub) status was assessed using Alexa Fluor 594-conjuagted αUb antibody. Images represent *Rickettsia*-infected macrophages after 2 h post-infection. (**B**) Graph shows the numbers of ubiquitin (Ub) positive stained *Rickettsia* spp per host cell after 2, 24 and 48 h post-infection. DNA was stained using DAPI (blue). Colocalization between *Rickettsia* and Ub was analyzed using Coloc 2 plugin Fiji software. Bars in panel A, 10 μm. (**C**) *Rickettsia*-infected WT BMDMΦ, as described in panel A, were lysed and samples were immunoblotted with αp62, αLC3B, αET-Ts, and αGAPDH Abs. Immunoblot data are a representative of three independent experiments. (**D**) Densitometry analysis from samples shown in (C) was performed using Fiji software and data represent the fold change between the ratios of LC3B-II/GAPDH, LC3B-II/LC3B-I or p62/GAPDH. Error bars (B, D) represent means ± SEM from five independent experiments; NS: non-significant; \*\**p* ≤ 0.01; \*\*\**p* ≤ 0.005; \*\*\*\**p* ≤ 0.001.

### Pathogenic *Rickettsia* species avoid autophagolysosomal destruction to establish a replicative niche in macrophages

Autophagy is an intracellular process that delivers autophagosome to the lysosome for degradation and is considered as one of host defenses to combat bacterial infection, including other cellular functions (18,19,38). To test our hypothesis that pathogenic, but not non-pathogenic, *Rickettsia* spp modulate autophagy flux, we evaluated the status of autophagy marker LC3B and lysosomal marker Lamp2, during infection of WT BMDMΦ by immunofluorescence assay (IFA). We observed that, unlike *R. montanensis, R. typhi*, and *R. rickettsii* colocalized with autophagy marker LC3B over the course of infection (**Figures 2A-D**). Given our preceding findings that *R. typhi* was able to avoid autophagolysosomal destruction in non-phagocytic cells (29), we evaluated the colocalization pattern of all three *Rickettsia* spp with Lamp2 as well as bacterial burdens. We observed that, unlike *R. montanensis*, both *R. typhi*, and *R. rickettsii* spp, did not colocalize with Lamp2 in WT BMDMΦ (**Figures 2A-E**), suggesting that highly pathogenic *Rickettsia* spp likely evade autophagolysosomal destruction to support intracytosolic replication.

**Figure. 2.**
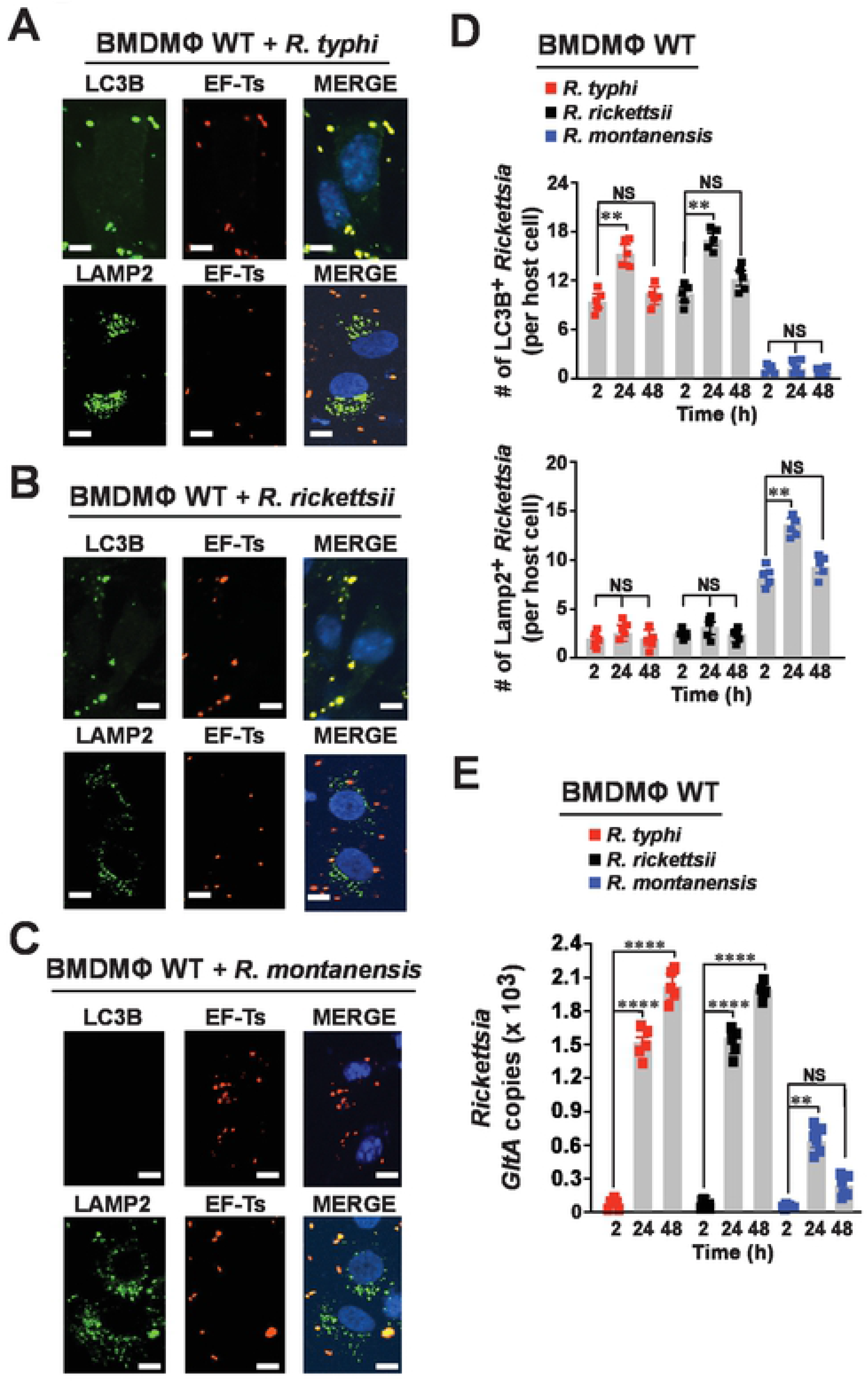
Pathogenic *Rickettsia* species evade autophagolysosomal destruction to establish a replication niche in macrophages. BMDMΦ from WT mice were infected with *R. typhi*, *R. rickettsii*, or *R. montanensis* for 2 h. (**A-C**) *Rickettsia*-infected BMDMΦ were fixed with 4% PFA and *Rickettsia* spp were detected using the rickettsial specific Alexa Fluor 594-conjugated EF-Ts, while LC3B, and Lamp2 expression was assessed using Alexa Fluor 488-conjuagted αLC3B, or αLamp2 antibodies respectively. Images represent *Rickettsia*-infected macrophages after 2 h post-infection. (**D**) Graphs show the numbers of LC3B, or Lamp2 positive stained *Rickettsia* spp per host cell after 2, 24 and 48 h post-infection. DNA was stained using DAPI (blue). Colocalization between *Rickettsia* and LC3B or Lamp2 was analyzed using Coloc 2 plugin Fiji software. Bars in panels A-C, 10 μm. (**E**) Bacterial burdens in infected BMDMΦ were evaluated 2, 24 and 48 h post-infection by *GltA* RT-qPCR. Expression of the host housekeeping gene *Gapdh* was used for normalization. Error bars (D-E) represent means ± SEM from five independent experiments; \*\**p* ≤ 0.01; \*\*\*\**p* ≤ 0.001.

### Evasion of autophagolysosomal destruction and reduction of anti-microbicidal pro-inflammatory IL-1 responses by pathogenic *Rickettsia* species involves an ATG5-dependent autophagic process

Based on recent findings from other laboratories and ours (21–24,29,31–34,39), it has become evident that *Rickettsia* spp exhibit variable pathogenicity, indicating species-specific strategies to respond to mechanism of host defense surveillance, including autophagy and inflammasome responses. In our efforts to address the role of autophagy in modulating the intracellular survival of *Rickettsia* spp, we first determined the colocalization status of *R. montanensis, R. typhi*, and *R. rickettsii* with LC3B during infection of ATG5^fl/fl^ or ATG5^fl/fl^-LysM-Cre BMDMΦ by IFA. We found that, unlike *R. montanensis*, both *R. typhi*, and *R. rickettsii* strains colocalized with LC3B in ATG5^fl/fl^ BMDMΦ, but not in ATG5^fl/fl^-LysM-Cre BMDMΦ (**Figures 3A-D**). Moreover, we observed that *R. montanensis*, but not *R. typhi* and *R. rickettsii*, colocalized with Lamp2 in ATG5^fl/fl^ BMDMΦ (**Figures 3A-D**), while all three *Rickettsia* strains colocalized with Lamp2 in ATG5^fl/fl^-LysM-Cre BMDMΦ (**Figures 3A-D**), suggesting that pathogenic *Rickettsia* spp avoid autophagosomal maturation in an ATG5-dependent manner (**Figures 3E**).

**Figure. 3.**
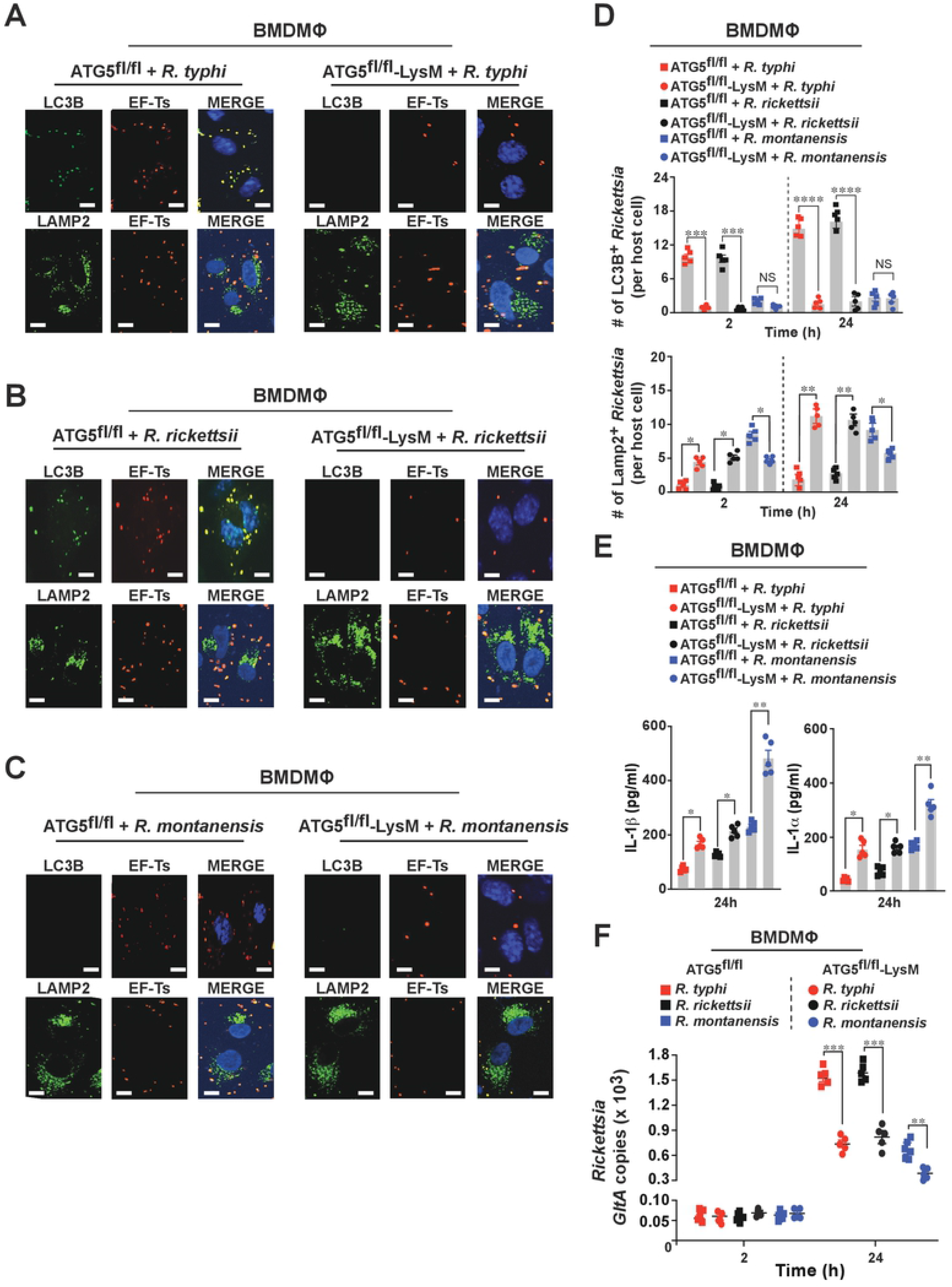
Evasion of autophagolysosomal destruction and suppression of anti-rickettsial inflammasome-dependent IL-1 cytokine responses by pathogenic *Rickettsia* involves an ATG5-mediated autophagic process. BMDMΦ from ATG5^fl/fl^ or ATG5^fl/fl^-LysM-Cre mice were infected with *R. typhi, R. rickettsii*, or *R. montanensis* for 2 h. (**A-C**) *Rickettsia*-infected ATG5^fl/fl^ and ATG5-deficient BMDMΦ were fixed with 4% PFA and *Rickettsia* spp were detected using the rickettsial specific Alexa Fluor 594-conjugated EF-Ts, while LC3B, and Lamp2 expression was assessed using Alexa Fluor 488-conjuagted αLC3B, or αLamp2 antibodies respectively. Images represent *Rickettsia*-infected macrophages after 2 h post-infection. (**D**) Graphs show the numbers of LC3B, or Lamp2 positive stained *Rickettsia* spp per host cell after 2 and 24 h post-infection. DNA was stained using DAPI (blue). Colocalization between *Rickettsia* and LC3B or Lamp2 was analyzed using Coloc 2 plugin Fiji software. Bars in panels A-C, 10 μm. (**E**) Culture supernatants of *Rickettsia*-infected ATG5^fl/fl^ and ATG5-deficient BMDMΦ were analyzed 24 h post-infection to determine the level of IL-1β and IL-1α cytokines using Legendplex kits (BioLegend) followed by flow cytometry. (**F**) Bacterial burdens in infected BMDMΦ were evaluated 2 and 24 h post-infection by *GltA* RT-qPCR. Expression of the host housekeeping gene *Gapdh* was used for normalization. Error bars (D-F) represent means ± SEM from five independent experiments; NS: non-significant; \**p* ≤ 0.05; \*\**p* ≤ 0.01; \*\*\**p* ≤ 0.005; \*\*\*\**p* ≤ 0.001.

Our recent findings identified a previously unappreciated mechanism by which highly pathogenic *Rickettsia* spp (*R. typhi*, and *R. rickettsii*), unlike non-pathogenic rickettsiae, benefitted from a reduced IL-1 signaling response, specifically IL-1α, to support their replication within the host (31). Intriguingly, *R. australis* benefited from ATG5-dependent autophagy induction and reduction of pro-inflammatory cytokine responses (21,22), however, a successful host colonization of *R. parkeri* involved the evasion of autophagy and inflammasome-mediated host cell death (23,24), leaving the precise mechanism inconclusive. To address this knowledge gaps, we tested the hypothesis that pathogenic *Rickettsia* spp reduce IL-1 cytokine responses via an autophagy-driven mechanism. In this effort, we measured the IL-1β and IL-1α cytokine levels in cultured supernatants of *R. montanensis-, R. typhi-*, and *R. rickettsii*-infected ATG5^fl/fl^ BMDMΦ, as well as ATG5^fl/fl^-LysM-Cre BMDMΦ. We observed that lack of autophagy was associated with an increase in the secretion of IL-1β or IL-1α upon infection with non-pathogenic as well as pathogenic *Rickettsia* spp (**Figure 3E**). In addition, bacterial burden assessments provided additional evidence that *Rickettsia* survival in macrophages is dependent on the function of ATG5 (**Figure 3F**). Collectively, our findings suggest replication of both pathogenic and non-pathogenic *Rickettsia* spp in BMDMΦ requires a ATG5-dependent autophagic mechanism.

Given these data, we sought to study further the biological importance of autophagy induction for pathogenic and non-pathogenic *Rickettsia* spp by utilizing a pharmacological inhibitor of autophagy, 3-methyladenine (3-MA), as well as an activator of autophagy, rapamycin (RA). Similar to our infection studies using ATG5-deficient BMDMΦ, treatment of WT BMDMΦ with 3-MA significantly decreased the number of LC3B^+^-bacteria as compared to levels observed in non-treated WT BMDMΦ (**Figures 4A-F, J, 24h**). Importantly, 3-MA treatment increased the number of Lamp2^+^-pathogenic *Rickettsia* (**Figures 4A-F, J**), supporting to our hypothesis that pathogenic *Rickettsia* spp rely on the initiation of autophagy and escape from autophagolysosomal destruction to establish a replication niche. On the other hand, autophagy induction in WT BMDMΦ via RA treatment significantly increased the number of LC3B^+^-bacteria of pathogenic and non-pathogenic *Rickettsia* spp (**Figures 4G-J, 24h**). Strikingly, RA treatment resulted in a significant decrease in the number of Lamp2^+^-*R. montanensis* bacteria (**Figures 4G-J**). Furthermore, examination of *Rickettsia* burdens in 3-MA and RA-treated WT BMDMΦ further highlighted the importance of autophagy initiation for pathogenic *Rickettsia* spp survival in macrophages.

**Figure. 4.**
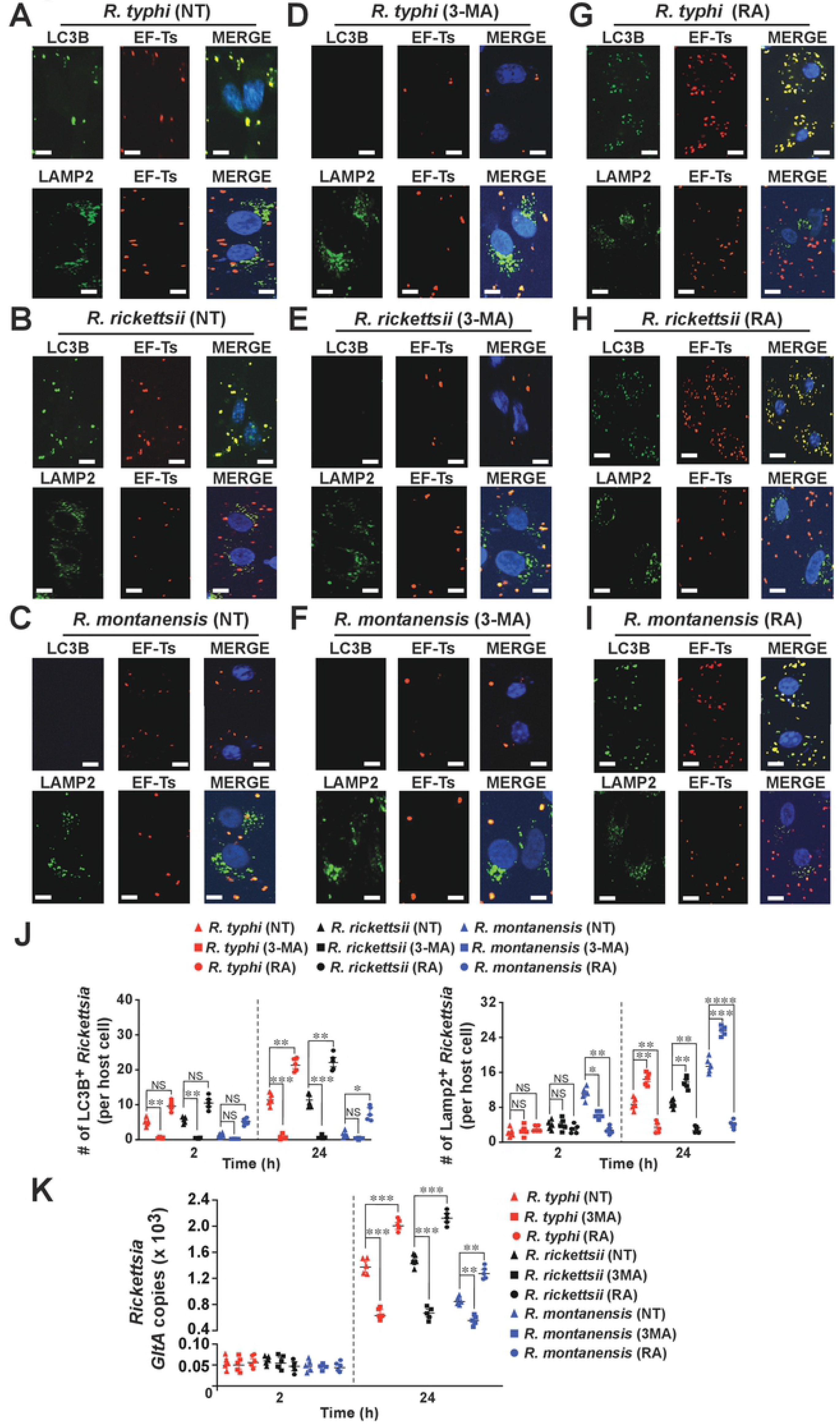
Pharmacological modulation of autophagy underlines the importance of autophagy to limit *Rickettsia* invasion in macrophages. (**A-I**) BMDMΦ from WT mice were pretreated with 3-methyladenine (3-MA, 5 mM) or rapamycin (RA, 0.1 mM) for 1 h followed by *R. montanensis, R. rickettsii*, or *R. typhi* infection for additional 2 h. Samples were fixed with 4% PFA and *Rickettsia* spp were detected using the rickettsial specific Alexa Fluor 594-conjugated EF-Ts, while LC3B, and Lamp2 expression was assessed using Alexa Fluor 488-conjuagted αLC3B, or αLamp2 antibodies respectively. (**J**) Graphs show the numbers of LC3B, or Lamp2 positive stained *Rickettsia* spp per host cell after pretreated with 3-MA or RA (2 and 24h). engulfment of cells in which *Rickettsia* spp colocalized with LC3B, or Lamp2. DNA was stained using DAPI (blue). Colocalization between *Rickettsia* and LC3B or Lamp2 was analyzed using Coloc 2 plugin Fiji software. Bars in panels A-I, 10 μm. Error bars represent means ± SEM from five independent experiments; NS: non-significant; \**p* ≤ 0.05; \*\**p* ≤ 0.01; \*\*\**p* ≤ 0.005; \*\*\*\**p* ≤ 0.001.

Collectively, our data are in agreement with previous reports (21,22,29,31), indicating that, unlike non-pathogenic *R. montanensis* spp, both pathogenic *R. typhi*, and *R. rickettsii* spp, induce autophagy, but evade autophagolysosomal destruction, and reduce anti-rickettsial inflammasome-dependent IL-1 cytokine responses to establish an intracytosolic replication niche in macrophages.

## Discussion

Various intracellular bacterial pathogens have developed sophisticated mechanisms to hijack host cellular processes to facilitate their survival. Such strategies entail reprogramming host phosphoinositide (PI) metabolism, which can facilitate uptake into host cells, modify phagosomes, undermine apoptosis, and interfere with other cellular defense mechanisms, such as inflammasomes and autophagy. However, in the case of *Rickettsia* the mechanisms by which these pathogens modulate both inflammasome and autophagy responses to facilitate their replication in endothelial cells and immune cells, like macrophages, is only now emerging (21,22,40,23,24,29,31–34,39). In fact, recent findings from others and our laboratory have provided some insight on how highly pathogenic *Rickettsia* spp modulate inflammasome-mediated immune responses to establish an intracellular niche. However, the precise role of autophagy in regulating host colonization by *Rickettsia* spp remains to be determined (21–24). For instance, *R. australis* benefited from autophagy induction and reduction of pro-inflammatory cytokine responses (21,22), while *R. parkeri* evaded autophagic responses to colonize the host (23,24). Intriguingly, our data on *R. typhi*, showed that ubiquitination followed by autophagy induction, but the escape from autophagosomes were crucial steps for *R. typhi* to colonize non-phagocytic cells (29). Based on these unexpected findings, we tested the hypothesis that highly pathogenic, but not non-pathogenic, *Rickettsia* spp induce autophagy, but evaded autophagolysosomal destruction, and reduces inflammasome-mediated anti-microbicidal pro-inflammatory IL-1 responses to establish a replication niche in host immune defense cells, like macrophages.

Our data revealed that, like *R. parkeri, R. montanensis* is not ubiquitinated during infection of macrophages, while both *R. typhi*, and *R. rickettsii* are ubiquitinated upon macrophages entry, which are in agreement with reports on *R. australis* (22). In addition, we observed that, unlike *R. montanensis, R. typhi*, and *R. rickettsii* colocalized with LC3B, but not with the lysosomal marker Lamp2, during macrophage invasion. Importantly, our presented data using the pharmacological modulators of autophagy (3-MA and RA), or ATG5-deficient macrophages further strengthen our hypothesis that pathogenic, but not non-pathogenic, *Rickettsia* spp induce autophagy and avoid autophagolysosomal maturation to establish an intracytosolic replication niche in macrophages (**Figure 5**).

**Figure. 5.**
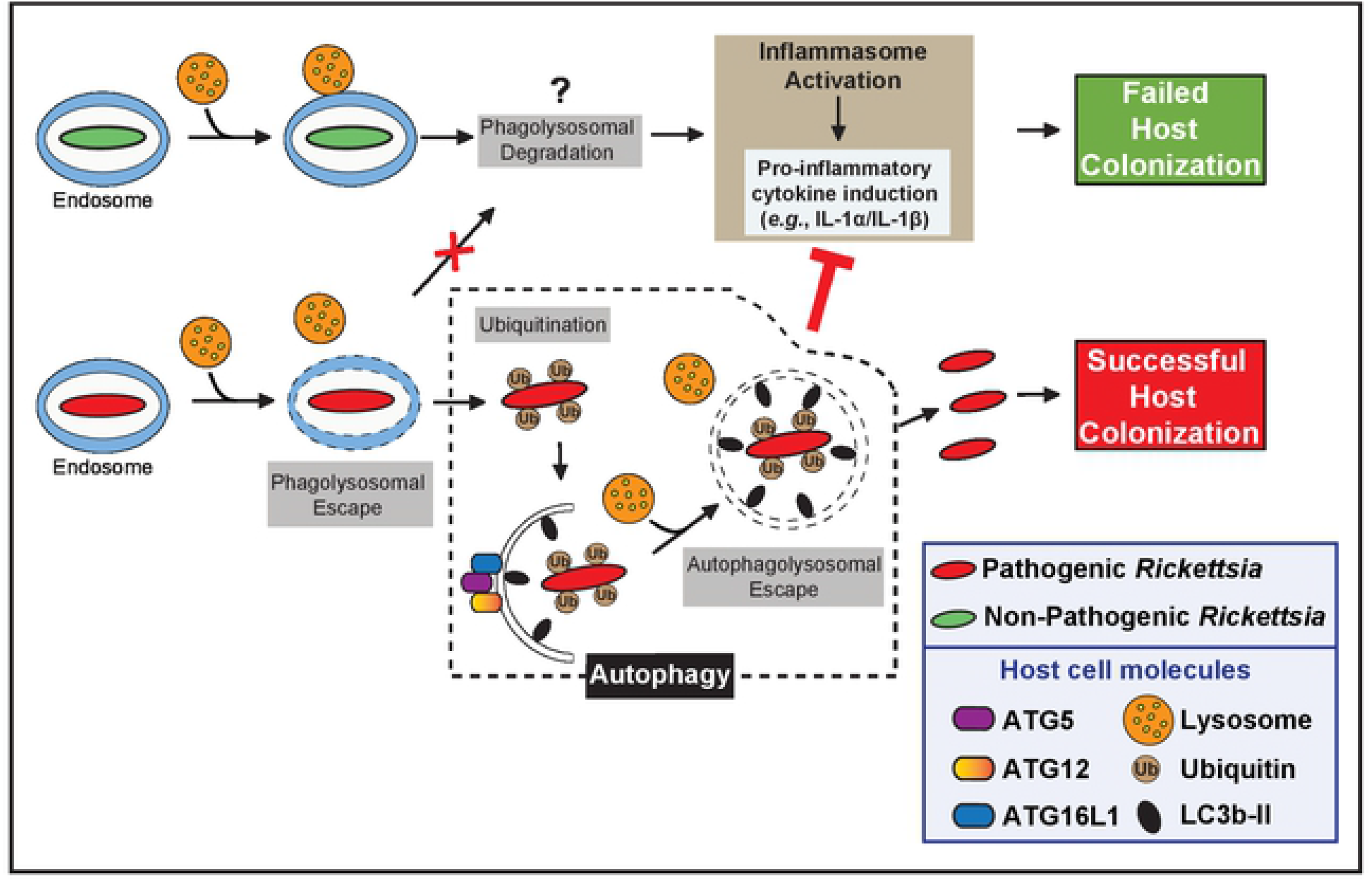
Proposed infection model of pathogenic *Rickettsia* species in macrophages. Proposed working model on how pathogenic *Rickettsia* spp initiate autophagy, evade autophagolysosomal destruction, and suppress anti-rickettsial inflammasome-dependent IL-1 cytokine responses to establish a replication niche in macrophages. Of note, the majority of non-pathogenic *Rickettsia* spp is likely destroyed by phagolysosomal fusion, while a subpopulation may escape lysosomal fusion, ultimately resulting in the induction of IL-1 responses.

Our findings also raised another intriguing question as to why *R. typhi*, and *R. rickettsii* spp seem to possess the ability to actively induce autophagy, while *R. montanensis* did not. As *Rickettsia-specific* immunodominant outer membrane proteins (*e.g*., Scas) are predicted to be expressed on either pathogenic or non-pathogenic *Rickettsia* spp (41), one alternative explanation is the presence of an effector repertoire that differs depending on the rickettsial virulence. In fact, we recently reported that *R. typhi* induces autophagy upon infection of non-phagocytic cells, while subsequently avoiding autophagosomal destruction, via the function of secreted effectors, including phosphatidylinositol 3-kinase Risk1 and phospholipases Pat-1 and −2 (29,41). This mechanism of host invasion seems consistent with rickettsial close relatives, such as *Anaplasma phagocytophilum* and *Ehrlichia chaffeensis* (42–45), or other intracellular pathogens such as *Shigella* (46,47). So, it is tempting to speculate that effectors [e.g., Risk1 and Pat1/2 (29,41)], either alone, or in combination with other currently unknown effectors, could account for the observed phenotypic differences in host colonization by pathogenic and non-pathogenic *Rickettsia* spp. The precise mechanism and composition of the effector repertoire for each *Rickettsia* spp is currently a matter of active research and remains to be determined.

Our preceding report suggest that IL-1 signaling responses played a role in limiting rickettsial infection *in vivo* and *in vitro* (31). Specifically, we observed that *R. typhi*, and *R. rickettsii* spp, but not *R. montanensis*, reduces non-canonical inflammasome-dependent IL-1α responses, in order, to establish an intracytosolic replication niche in macrophages. In fact, we reported that non-pathogenic, but not pathogenic, *Rickettsia* spp are more efficiently cleared by macrophages, a mechanism supported the previously published findings with SFG *Rickettsia* using THP-1 cells (48). Given our recent reporting (29,31) and by others (13,14,17,18), we evaluated the IL-1 cytokine responses and bacterial burden in ATG5-deficient macrophages and observed ATG5-mediated autophagic processes played a key role in regulating IL-1 responses that limit the replication of both pathogenic and non-pathogenic *Rickettsia* spp in macrophages, supporting the pathway by which autophagy acts as a negative regulator of inflammasome responses (17,49).

Overall, our findings present a working model of host invasion by which pathogenic, but not non-pathogenic, *Rickettsia* spp become ubiquitinated, induced autophagy, but avoid autophagolysosomal destruction, and reduces inflammasome-mediated anti-microbicidal IL-1 cytokine responses to establish an intracytosolic niche in macrophages (**Figure 5**).

## Materials and Methods

### Antibodies and Reagents

Mono- and polyubiquitinylated conjugated antibody [clone: FK2] (αUb) was obtained from Enzo Life Sciences. Anti-LC3B (E5Q2K) antibody was from Cell Signaling Technology. Antibody against rickettsial cytoplasmic housekeeping protein elongation factor Ts (EF-Ts) was obtained from Primm Biotech as previously described (29). The p62/SQSTM1 antibody, 3-methyladenine (3-MA), and rapamycin (RA) were purchased from Sigma. Lamp2 (H4B4) and GAPDH (FL-335) antibodies were from Santa Cruz Biotechnology. ProLong Gold antifade mounting medium with DAPI (4’,6-diamidino-2-phenylindole), Halt protease and phosphatase inhibitor cocktail, paraformaldehyde (PFA), and Alexa 488/594-conjugated secondary Abs were purchased from Thermo Fisher Scientific.

### Bacterial strains, cell culture, and infection

Vero76 cells (African green monkey kidney, ATCC, RL-1587) were maintained in minimal Dulbecco’s modified Eagle’s medium (DMEM) supplemented with 10% heat-inactivated fetal bovine serum (FBS) at 37°C with 5% CO_2_. *R. montanensis* strain M5/6 and *R. rickettsii* (Sheila Smith) strain were obtained from Dr. Ted Hackstadt (Rocky Mountain Laboratories, NIH, MT, USA) and *R. typhi* strain Wilmington was obtained from CDC. All *Rickettsia* strains were propagated in Vero76 cells grown in DMEM supplemented with 5% FBS at 34°C with 5% CO_2_. All *Rickettsia* were purified as previously described (26). For infection of BMDMΦ, purified *Rickettsia* spp were used at a multiplicity of infection (MOI) of 50-100, to ensure the presence of enough bacteria at early stage of infection, for host responses (25–27,29,31).

### Differentiation of bone marrow-derived macrophages

Bone marrow cells were isolated from femurs and tibias of WT, ATG5^fl/fl^, and ATG5^fl/fl^-LysM-Cre mice. Femurs from ATG5^fl/fl^ and ATG5^fl/fl^-LysM-Cre mice were provided by Dr. Christina Stallings (Washington University School of Medicine, MO, USA). Differentiation was induced by culturing bone marrow cells in RPMI 1640 medium supplemented with 10% FBS and 30% L929-conditioned medium (a source of macrophage colony stimulating factor) and cultured for 7 days as described previously (31,50).

### Measurement of cytokines and chemokines

IL-1 cytokine concentrations in supernatants from cultured BMDMΦ were assessed using the Legendplex mouse inflammation kit (BioLegend) following the manufacturer’s instructions as described previously (31,50).

### RNA isolation and quantitative real-time PCR

To determine viable bacterial number during the course of host infection, we performed RT-qPCR assay on isolated RNA (25,51,52). In this effort, BMDMΦ samples were collected at 2, and 24h post-infection. RNA was extracted from 1 × 10^6^ BMDMΦ using the Quick-RNA miniprep kit (ZymoResearch). The iScript Reverse Transcription Supermix kit (Bio-Rad; 1708841) was used to synthesize cDNAs from 200 ng of RNA according to the manufacturer’s instructions. Quantitative real-time PCR was performed using SYBR Green (Thermo Fisher Scientific), 2 μl cDNA and 1 μM each of the following oligonucleotides for rickettsial citrate synthase gene (*gltA*), and host gene *Gapdh*. Relative rickettsial citrate synthase gene (*gltA*) expression was measured using the comparative threshold cycle (2^−ΔΔCT^) method with host gene *Gapdh* as the reference transcript to normalize the bacterial burdens in host cells as described previously (31,53).

### Immunofluorescence

Eight-well chamber slides were seeded with BMDMΦ (30-50 × 10^4^ cells/well) and infected with partially purified pathogenic and non-pathogenic *Rickettsia* spp (MOI 50-100) as described previously (25–27,29,31). Briefly, partially purified *Rickettsia* were added to BMDMΦ and incubated for various length of time at 34°C with 5% CO_2_. Following incubation, cells were washed three times with 1 x PBS and fixed with 4% paraformaldehyde (PFA) for 20 min at room temperature. Cells were than permeabilized in blocking buffer (0.3% saponin and 0.5% normal goat serum in 1 x PBS) for 30 min and incubated for 1 h with the following primary Abs diluted in Ab dilution buffer (0.3% saponin in 1 x PBS): αEF-Ts (1:500), αUb (1:100), αLC3B (1:100), and αLamp2 (1:100). Cells were then washed with 1 x PBS and incubated for 1 h with αAlexa Fluor 488 or αAlexa Fluor 594 secondary Abs diluted 1:1,500 in Ab dilution buffer. Next, cells were washed with 1 x PBS and mounted with ProLong Gold antifade mounting medium containing DAPI. Images were acquired using the Nikon W-1 spinning disk confocal microscope (University of Maryland Baltimore, Confocal Core Facility) and degree of colocalization (yellow^+^-stained bacteria) between *Rickettsia* and ubiquitin, LC3B or Lamp2 was analyzed using Fiji software as described previously (29). Approximately 100 host cells were enumerated for each experiment and each experiment was repeated five times.

### Extract preparation, and Western blot analysis

*Rickettsia*-infected BMDMΦ cells were lysed for 2 h at 4°C in ice-cold lysis buffer (50 mM HEPES [pH 7.4], 137 mM NaCl, 10% glycerol, 1 mM EDTA, 0.5% NP-40, and supplemented with protease and phosphatase inhibitory cocktails) as described previously (29,31). Equal amounts of protein were loaded for SDS-PAGE and membranes were probed with αp62, αLC3B, αEF-Ts, and αGAPDH Abs, followed by enhanced chemiluminescence with secondary Abs conjugated to horseradish peroxidase.

### Statistical analysis

The statistical significance was assessed using ANOVA with Bonferroni’s procedure and Student t-test (GraphPad Prism Software, version 8). Data are presented as mean ± SEM, unless stated otherwise. Alpha level was set to 0.05.

## Acknowledgments

We gratefully acknowledge Christina Stallings (Washington University School of Medicine, MO, USA), and Ted Hackstadt (Rocky Mountain Laboratories, NIH, MT, USA) for generously providing us with essential biological specimens and reagents, including femurs from various knock-out mice and rickettsial strains. We would like to thank Magda Beier-Sexton for her administrative, technical, organizational, and editorial contributions to the manuscript. This work was supported with funds from the NIAID/NIH grants (R01AI017828 and R01AI126853 to A.F.A.).

